# Modeling Protein-based Hydrogels under Force

**DOI:** 10.1101/361592

**Authors:** Kirill Shmilovich, Ionel Popa

## Abstract

Hydrogels made from structured polyprotein domains combine the properties of cross-linked polymers with the unfolding phase transition. The use of protein hydrogels as an ensemble approach to study the physics of domain unfolding is limited by the lack of scaling tools and by the complexity of the system. Here we propose a model to describe the biomechanical response of protein hydrogels based on the unfolding and extension of protein domains under force. Our model takes into account the contributions on the network dynamics of the molecules inside the gels, which have random cross-linking points and random topology. This model reproduces reported macroscopic visco-elastic effects and constitutes an important step toward using rheometry on protein hydrogels to scale down to the average mechanical response of protein molecules.

---

Protein-based hydrogels are a new type of material that retains the main characteristics of polymeric hydrogels, but shows a unique visco-elastic response to stress. This response stems from the unfolding and extension of constituent protein domains. The appearance of the unfolding phase transition depends on the experienced force, exposure time, pulling geometry, and what protein is used to form the gel. Such a unique response of protein-based hydrogels to external stimuli does not only open new vistas toward designing new biological materials, but also enables a new spectroscopy technique to determine the mechanical response and energy landscape of single proteins from multi-molecule ensemble experiments of protein hydrogels. Rather than gathering single-molecule data through time-consuming atomic force microscopy (AFM) or optical and magnetic tweezers measurements, soft-matter rheometry can probe the force response of a massive number of interconnected proteins [1,2]. Rheometry techniques require the decoupling the force-induced (un)folding of individual proteins from the elastic response related to the cross-linked gel network, an experiment that recently became available through the introduction of force-clamp rheometry [3].

Here we propose a model that describes the macroscopic response of protein-based hydrogels obtained from polyproteins. This model is a critical step toward extracting the average unfolding and extension of single molecules from hydrogel stretching experiments.

Polyproteins have a cylindrical geometry and can be cross-linked into hydrogels using a photo-activated chemical reaction, where exposed tyrosine amino acids produce carbon-carbon bonds between adjacent polyprotein molecules [1]. Once the protein network is formed, its response to force can be analyzed through the dynamics of its cross-linking nodes. An elegant approach was introduced to model the network dynamics of actin filaments under a perturbing force vector [4-6]. While actin domains do not experience any unfolding or refolding transitions, the cylinder like geometry of actin filaments resembles that of polyproteins.

Unique to polyprotein hydrogels is the unfolding transition of constituent domains, which results in a significant increase in the contour length of the molecule. We chose to investigate hydrogels made from polyproteins (repeats of protein L), as this model protein has been extensively studied experimentally by the single molecule force spectroscopy community [2,7]. The domains in polyproteins are arranged as ‘beads on a string’. This arrangement is an important characteristic of many proteins that have evolved to operate under force, such as titin in muscles [8] and talin in cellular mechano-transduction [9,10]. Furthermore, an energy landscape model for a polyprotein made of eight repeats of protein L was shown to reproduce the measured unfolding and refolding response of this protein to force, and was adopted herein [11,12]. This model combines the change in the barrier height between the folded and unfolded states due to an applied force with standard polymer elasticity models, which account for the entropic extension of the unfolded polypeptide chain.

As hydrogels are over 90% water, it is reasonable to assume that the dynamics of individual molecules inside hydrogels is the same as the dynamics of the polyprotein measured by force spectroscopy in solution. Our model ignores any intermediate states that characterize the folding process, as these are short-lived [13] when compared to the chosen sampling time. Furthermore, as most of the domains composing a molecule will not be part of the cross-link, it is safe to assume that they will experience the force along the polyprotein N-to-C backbone, which is the same pulling coordinate as in single molecule experiments. Those domains that are part of the crosslinks will have a different stability [14], but their overall effect on the gel dynamics is limited, as they only partially extend between the cross-link and either the N or C-terminus (see also below equation (4)).

In our model, each polyprotein L molecule is approximated to a rigid rod, with *m* = 8 domains of radius *r* = 2 *nm* each (PDB code *1hz5*), leading to a total contour length *L* = 32 *nm.* To form the gel network, each polyprotein molecule was assigned a center of mass, following a random distribution inside a square lattice, along with a random orientation. *N* polyprotein molecules were distributed within a volume *V* = *N/c*, where *c* is the molecule number density. The proteins were allowed to diffuse inside a rigid box of a volume *3V* with a mean square displacement 〈*x*^2^〉 = 2*D_t_*δ*t* and to rotate with 〈*φ*^2^〉 = 2*D_r_δt*, while interacting elastically with the box walls [15,16]. The translation and rotation diffusion coefficients for a single polyprotein molecule are defined as:

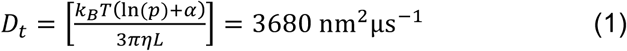

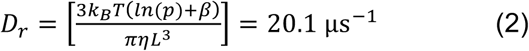

with *p* = *L*/(2*r*) being the shape factor for a rod, and α and β second degree polynomials in *p*^−1^[15,16]. Crosslinking occurs if the center of mass of adjacent protein domains are within a threshold distance 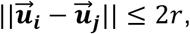 where 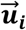 points to node *i* of the network.

When cross-linked, clusters of two or more molecules move in tandem with a diffusion coefficient of 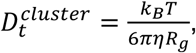 where *R_g_* is the radius of gyration [17]. Following complete crosslinking, the network was allowed to equilibrate using a quasi-Newton algorithm that shifts the position of cross-linking nodes to minimize

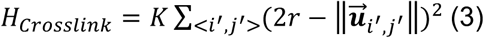

The primed indices restrict the summation to all valid inter-protein (i.e. cross-linked) node indices, with *K* = 3.72 · 10^5^ pN/nm being the force constant associated with the quadratic approximation of a C_n_-C_n_ bond [18], and 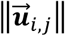 the perturbed bond length, such that 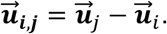

Under a constant force 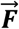 applied to the entire gel, the force experienced by a single molecule 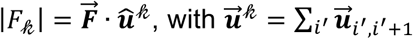 being determined by the nodes along molecule *𝓀* (Fig. 1A). Force can lead to unfolding and extension of protein domains along polyprotein molecules. An unfolding or refolding event of a domain *j* on polyprotein molecule *𝓀* extends/contracts the total end-to-end length *x_𝓀_* by an amount Δ along *û*^𝓀^.

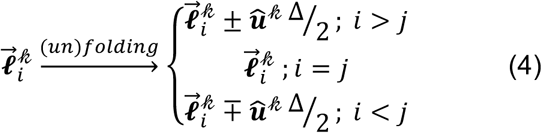

**Figure 1.**
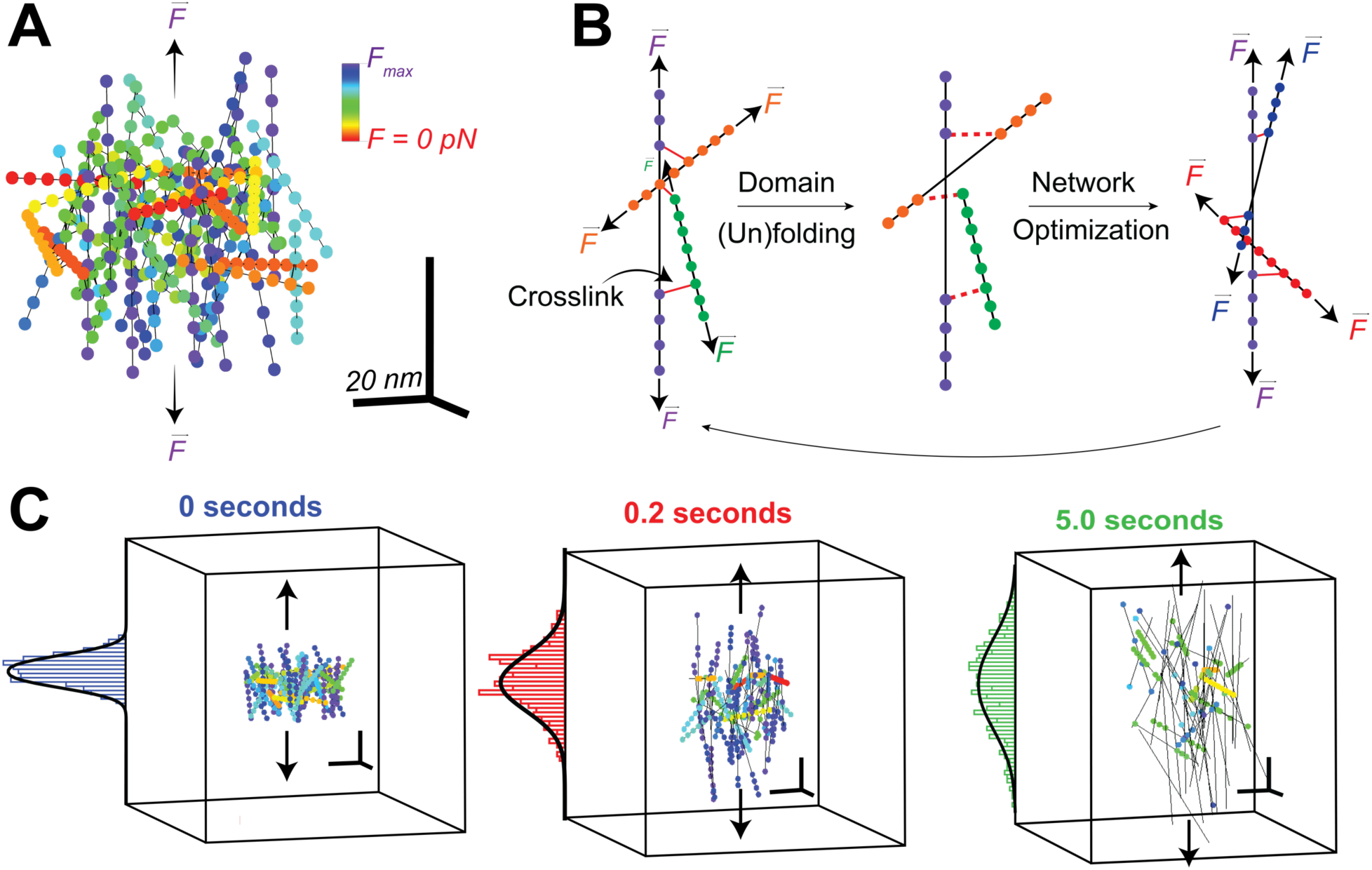
Protein hydrogels under force. (A) Molecules are depicted as straight lines with eight spheres along their axis which represent folded protein domains. A global constant force is applied to the z-axis. (B) Illustration of the dynamic unfolding and orientation correction during gel stretching. (C) Snap-shots of the same gel at three different time points. Scale bars are 20×20×20 nm.

Where ‘+’ signifies an unfolding event and ‘–’ a refolding transition. To understand how the (un)folding of single domains perturb the entire gel network, we adopt a formalism used to describe actin gels [6], which minimizes the stretching and bending terms:

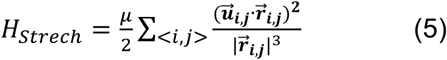

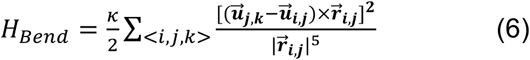

where 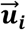 and 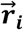 point to nodes in the perturbed and stable configurations respectively. The coefficients *κ* and *μ* are proportional to the persistence length *ℓ_p_*, and related to the geometry of the network constituents [19]. For a rigid uniform rod-like polyprotein, *κ* = *ℓ_p_k_B_T* = 2.4 pN · nm^2^ and 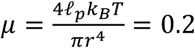 pN, where *ℓ_p_* = 0.58 nm for protein L [2]. Each network node is shifted during network optimization to minimize the total energy *H_Total_* = *H_Crosslink_* + *H_Bend_* + *H_Strech_* (Fig. 1B). This network optimization step was carried out only following unfolding or refolding events, as only then there is a significant change in the end-to-end length of a molecule inside the gel. The total gel extension was estimated by projecting all the molecules on the stretching coordinate *z*, and fitting a higher-order Gaussian function[20]: *F_P_*(*z*, *t*) = *A* exp[–(*z*/*γ*)^2*P*^], where 2*γ* represents the gel length (Fig. 1C).

Under constant force, polyproteins show probabilistic unfolding events in single molecule experiments, resulting in a stair-case like extension, rather than one large step at a well-defined time [2]. As more proteins participate in the overall mechanical response, this probabilistic response is expected to be smeared out. Indeed, when increasing the number of molecules that are used to form the hydrogel network, we observe a decrease in the variance between individual extension traces and the average for a given force protocol (Fig. 2). In this case, five separately polymerized hydrogels composed of *N* = 1, 8, 18, 32, 40, 50, 60, and 72 proteins at a concentration *c* = 9.0 · 10^−4^ molecules/nm^3^ (~1.5 mM) were exposed to a constant applied force of *F* = 50 pN for 5 seconds. The residual of the extension between the average and individual traces decreases with an increasing number of molecules, and stabilizes to ~4% for *N* ≥ 50. These results agree with the correspondence principle and tend toward the strictly deterministic behavior observed in tissues and biomaterials [21]. Indeed, rheometry measurements of protein hydrogels show very little change between measured elastic responses under identical conditions [3].

**Figure 2.**
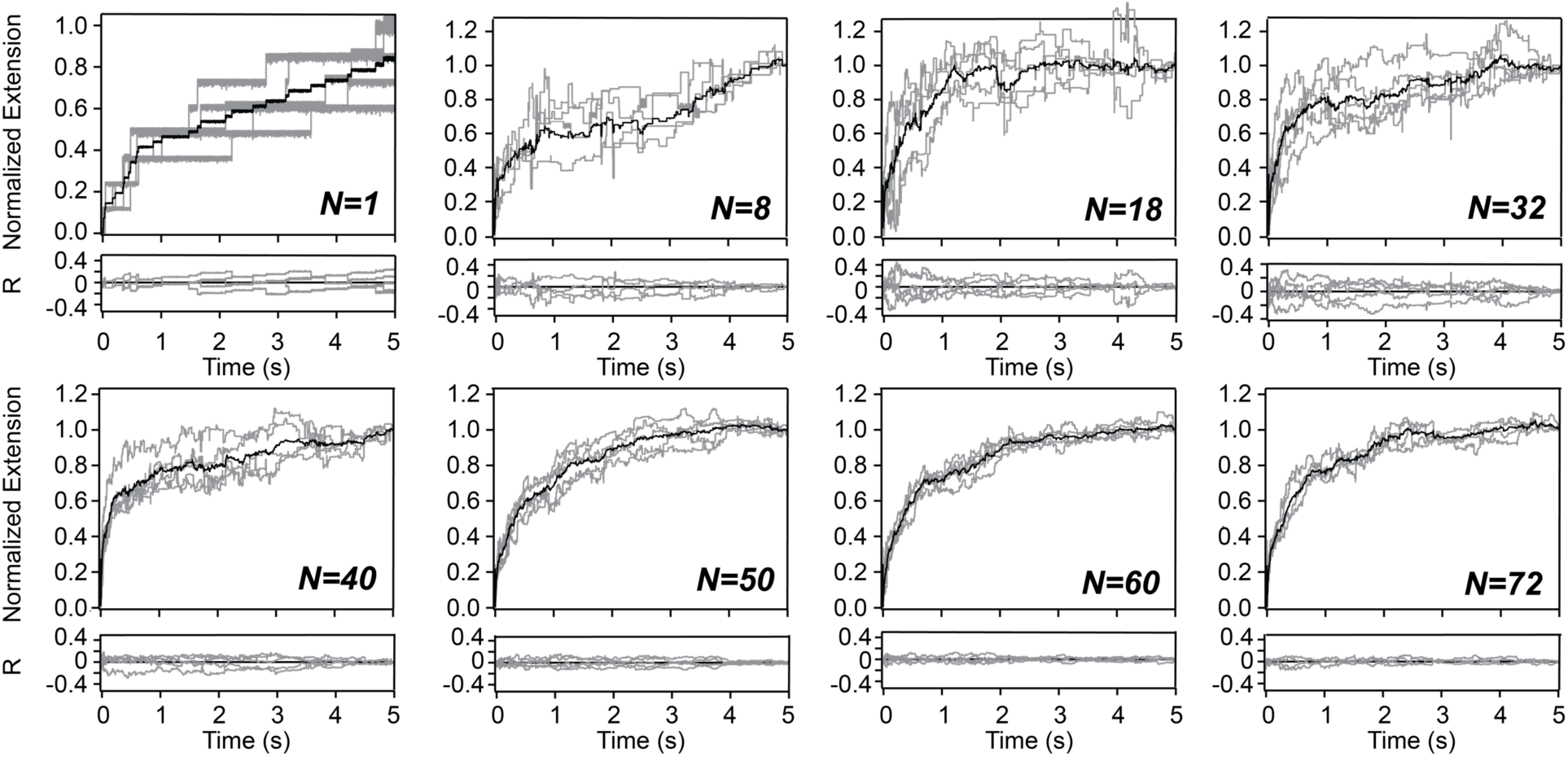
Scaling behavior of protein hydrogel extension at constant force. Hydrogels composed of varying number of proteins (from *N* = *1* to *N* = *72*) at a concentration of *c* = 9.0 · 10^−4^ molecules/nm^3^are subjected to an applied force of 50 pN for 5 seconds. Five separately polymerized hydrogels are simulated at each hydrogel size (grey traces) and averaged (black traces) with each trace being end-normalized. The residuals *R* between the average trace and the individual traces is shown below each graph.

To investigate the extension of hydrogels to mechanical forces using our model, we simulated networks made of *N* = 50 molecules at a concentration *c* = 9.0 · 10^−4^ molecules/nm^3^, which we find optimal in terms of probabilistic hydrogel response and computation time. Our simulations reproduce the measured behavior of protein hydrogels at constant force, which showed an initial elastic response, followed by a slower visco-elastic regime (Fig. 3A) [3]. As previously reported [22], a single exponential law describes poorly the visco-elastic regime. Complex visco-elastic materials with multiple mechanisms underlying their force response can be modeled using the Maxwell–Wiechert model [23]: a parallel assemblage of separately parameterized springs and dashpot Maxwell elements *x*(*t*) = Σ_*i*_*a_i_e*^−*τ_i_t*^, where *τ_i_* is the corresponding rate. Indeed, we, as well, find that a two-term exponential model fits our data best, but now we also have a clearer picture of what each exponential reports on. The fast rate constant is dominated by the initial alignment of molecules to the applied force and, at high forces, some unfolding events take place in molecules already aligned on the direction of the force vector (green triangles in Fig. 3B). The slow rate constant, on the other hand, are dominated by individual protein unfolding events and sporadically occurring rearrangements of the network have a relatively small effect (blue squares in Fig. 3B). Interestingly, for forces below ~55 pN force-per-molecule, the slower unfolding rates of hydrogels remain higher than the single-molecule rates (red circles), as the contribution from the network dynamics also contribute to this response.

**Figure 3.**
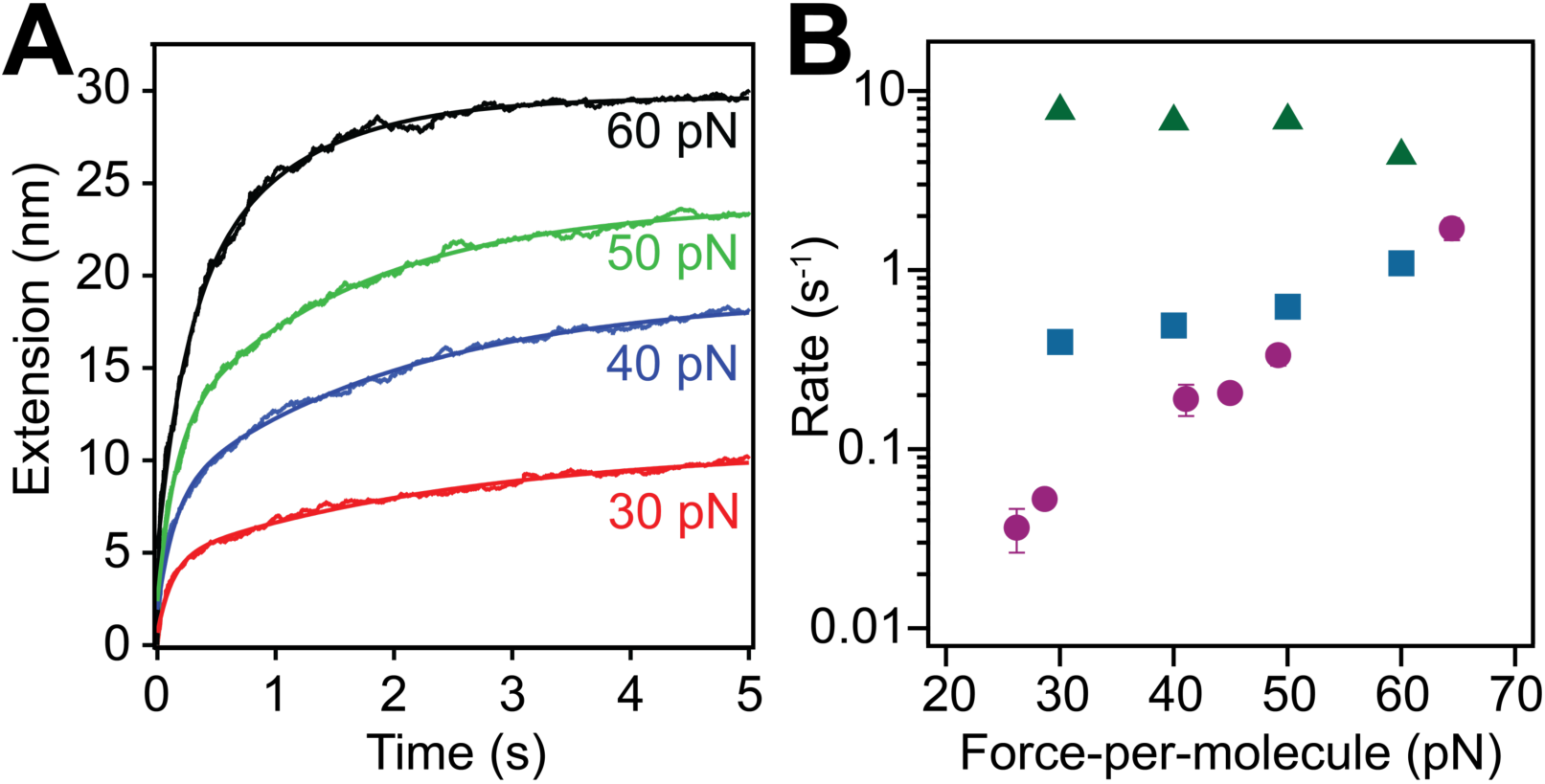
Force dependent behavior of protein-based hydrogel extension. (A) Average of 15 different hydrogels traces of *N* = 50 molecules at a concentration of *c* = 9.0 · 10^−4^ molecules/nm^3^, subjected to a constant force of 30, 40, 50, and 60 pN. (B) Comparison between rate constants of single molecule unfolding (circles) and rate constants from gels, determined by two-term exponential fits from (A) plotted as a function of the applied force. The faster, elasticity driven rates (triangles) stay relatively constant while the slower, unfolding driven rates (square) increase with force.

In summary, we propose a model to simulate the mechanical properties of protein-based hydrogels. This model builds on an established model which describes the unfolding response of polyproteins to a force along their end-to-end coordinate. Our model assumes no breaking of covalent bonds and utilizes the constraint imposed by domain cross-linking to optimize the network dynamics. The network equilibrates at the cross-linking points, following unfolding/refolding events. As the number of molecules forming a gel is increase, this model successfully recovers the probabilistic to deterministic scaling behavior expected in aggregating the stochastic process of polyprotein (un)folding. Furthermore, we have investigated the force-dependency of small protein-based hydrogels (*N* = *50*) to better understand the contributions from individual protein (un)folding events and the deformation mechanics of the crosslinked hydrogel network. Extension traces fit the multi-term exponential behavior commonly attributed to viscoelastic materials [23]. The faster rate is attributed to the initially quick elastic response of the network nodes, while the slower rate characterizes to the extended viscoelastic response. The relative ease of applying the presented formalism to hydrogels of generic protein composition furnishes an exciting new approach to probe the nanoscale behavior of protein-based hydrogels and offers a new way to extract unfolding and refolding dynamics from hydrogel rheometry measurements.

## Acknowledgments

This work was funded by Research Growth Initiative (Award No. 101X340), National Science Foundation, Major Research Instrumentation Program (Grant No. PHY-1126386), Greater Milwaukee Foundation (Shaw Award) and University of Wisconsin System (Applied Research Grant). K.S. also acknowledges funding from National Science Foundation (Grant No. UBM-1129056).

